# Zebrafish anterior segment mesenchyme progenitors are defined by function of tfap2a but not sox10

**DOI:** 10.1101/2022.10.20.513081

**Authors:** Oliver Vöcking, K Van Der Meulen, M.K Patel, J.K Famulski

**Affiliations:** Department of Biology, University of Kentucky

**Keywords:** anterior segment, neural crest, periocular mesenchyme, cornea, zebrafish, tfap2

## Abstract

The anterior segment is a critical component of the visual system. Developing independent of the retina, the AS relies partially on cranial neural crest cells (cNCC) as its earliest progenitors. The cNCCs are thought to first adopt a periocular mesenchyme (POM) fate and subsequently target to the AS upon formation of the rudimentary retina. AS targeted POM is termed anterior segment mesenchyme (ASM). However, it remains unknown when and how the switch from cNCC to POM or POM to ASM takes place. As such, we sought to visualize the timing of these transitions and identify the regulators of this process using the zebrafish embryo model. Using two color fluorescence *in situ* hybridization, we tracked cNCC and ASM target gene expression from 12-24hpf. In doing so, we identified a *tfap2a* and *foxc1a* co-expression at 16hpf, identifying the earliest ASM to arrive at the AS. Interestingly, expression of two other key regulators of NCC, *foxD3* and *sox10* was not associated with early ASM. Functional analysis of tfap2a, foxd3 and sox10 revealed that tfap2a and foxd3 are both critical regulators of ASM specification and AS formation while sox10 was dispensable for either specification or development of the AS. Using genetic knockout lines, we show that in the absence of tfap2a or foxD3 function ASM cells are not specified, and subsequently the AS is malformed. Conversely, sox10 genetic mutants or CRISPR Cas9 injected embryos displayed no defects in ASM specification, migration or the AS. Lastly, using transcriptomic analysis, we show that GFP+ cNCCs derived from Tg[*foxd3*:GFP] and Tg[*foxc1b*:GFP] share expression profiles consistent with ASM development whereas cNCCs isolated from Tg[*sox10*:GFP] exhibit expression profiles associated with vasculogenesis, muscle function and pigmentation. Taken together, we identify the earliest stage of anterior segment mesenchyme (ASM) specification to be approximately 16hpf and involve tfap2a/foxc1a positive cNCCs.

## INTRODUCTION

One of the great mysteries of the anterior segment lies in the complex dynamics of its beginnings. The exact molecular mechanisms and cellular relationships that allow for the development of anterior segment (AS) structures are poorly understood. Neural crest cells (NCCs) play a key role as the cell type of origin for periocular mesenchyme (POM) derivatives, the progenitors of the anterior segment mesenchyme (ASM) which in turn populate the AS (1). NCCs can be divided into smaller subpopulations with distinct genetic characteristics, differentiation abilities, migratory behaviors, and location of origin along the anterior-posterior axis of the neural tube. These groups consist of cranial, cardiac, vagal, and trunk subtypes (2-6).The most anterior population of NCCs are the cranial NCCs (cNCC). This population is one of the most extensively studied NCCs as it gives rise to the most diverse cell types and is associated with the most congenital defects. CNCCs differentiate into facial skeleton, facial cartilage, glia cells, neurons of the sensory ganglia, pharyngeal arches which in turn give rise to the jaw, connective tissues, bones of the semicircular canal, dental components, ocular muscles, and portions of the ocular anterior segment (3, 6-8). CNCCs typically originate in the sections of the neural tube corresponding to the forebrain, midbrain, and first pharyngeal arches. CNCCs are specified during NCC induction and migration through the gene regulatory network (GRN), a matrix of transcription factors required for cNCC fate determination. Genes turned on at the specification of cNCCs include *cMyc, Foxd3, Ets-1, Twist, Sox8/9/10, tfap2a* and *Snail1/2* (3, 6, 9, 10). While not all are activated simultaneously, these genes help to give NCCs their identity by maintaining a multipotent state, controlling proliferation, migration, and survival through regulation of cell adhesion and motility, and initiating the epidermal-mesenchymal-transition (EMT) and delamination (3, 4, 6, 9, 10).

It has been established how various NCCs, including cNCCs, are specified. However, the mechanism how cNCC transition into POM and subsequently into anterior segment mesenchyme (ASM) (i.e. POM cells which have committed to the AS fate and migrated to the AS space) remains unclear (1). Numerous studies have shown that an absence of cNCC leads to improper AS formation (11). However, the mechanism that specifies and targets cNCC or POM to become ASM remains unknown. POM cells are a migratory population, largely composed of cNCC derived cells. Though, they also include a small proportion of mesoderm cells, that populate the space immediately surrounding the developing eye and are essential for the proper development of the AS (11-16). POM cells can essentially be thought of as one step further specified than their cNCC progenitors: still highly migratory with the capability of differentiating into several different cell types but lacking the broad pluripotency of their cNCC counterparts. If cNCCs are multipotent stem cell-like cells contributing to numerous different cranial structures, than POM cells are slightly more refined, able to differentiate into several different cell and tissue types, but not as expansively as the cNCC. One of the tissue types derived from POM are the ASM. Any disruption in the specification or migration of POM or ASM cells can give rise to AS dysgenesis (ASD) phenotypes and disorders (12, 15, 17, 18). ASD is an umbrella term for maldevelopment of the AS including the cornea, lens, iridocorneal angle (ICA) and iris and includes several congenital conditions. We recently characterized the molecular and behavioral properties of the ASM (1). Our findings indicated that colonization of the AS by the ASM cells involves several independent populations of ASM likely originating from various cNCC subpopulations. Populations assessed in our assay included cells derived from foxc1b, foxd3, sox10, pitx2 and lmx1b.1 lineages. ASM from these various sources exhibit distinct migratory, colonization and transcriptomic profiles. Based on transcriptomic results, we concluded that there is significant variation in differentiation status of the ASM cells as they colonize the AS. Some of the subpopulations displayed markers associated with maturing AS structures whereas others displayed those of a more cNCC pluripotent state. The AS associated subpopulations were largely derived from foxc1b, lmx1b and pitx2 lineages, while sox10/foxd3 derived lineages displayed the cNCC like states. While our studies characterized ASM behavior and molecular signatures, we did not determine how and when cNCCs transition into POM and then subsequently into ASM.

In our current study we sought to visualize the timing of the transition between cNCC and POM as well as to characterize the mechanisms regulating this switch. We hypothesized that key regulators of cNCC differentiation tfap2a, foxd3 and sox10 are also potential candidates for modulating the switch between cNCC, POM and ASM as their expression in ASM persists up to and beyond the time AS colonization.

Tfap2a belongs to the tfap2 family of transcription factors which include paralogs *tfap2b, c, d* and are conserved between mammals and zebrafish. Tfap2a has been shown to be a critical component of the GRN and necessary for proper specification of the cNCCs. Tfap2a and tfap2b have also been shown to play roles in aspects of ocular development, including formation of the neural retina and anterior segment. Mice exhibiting NCC specific knockouts of tfap2a or tfap2b display characteristics of glaucoma and corneal deficiencies (19, 20). In humans, loss of tfap2a function has been associated with branchio-oculo-facial syndrome which includes incidence of cleft palate and craniofacial abnormalities.

FoxD3 is a vitally important gene of the NCC gene regulatory network, broadly expressed throughout the pre-migratory NC and is needed for the induction, survival, migration, and pluripotency of NCCs (21-24). FoxD3 is first activated during the neural crest specifier gene stage of NC development (3, 4, 6, 9, 10). FoxD3 in zebrafish is active at the neural plate border during gastrulation and continues to be expressed throughout the pre-migratory stages (23-25). Like other NC genes, FoxD3 remains active during the process of migration until further differentiation can occur and has been shown to be required for many, but not all, derivatives of the cNCCs (10, 22). In mice, FoxD3 has been shown to be required for the continued pluripotency of stem cells and homozygous KO mutants are embryonic lethal (21, 24). In zebrafish lacking FoxD3, craniofacial defects, cardiac abnormalities, and embryonic lethality are often observed (22, 24). In chick, FoxD3 overexpression results in the inability to differentiate NCCs (24). FoxD3 variation has also been identified in human ASD patients. Several families with Peter’s Anomaly or aniridia exhibit variations in the conserved region of FoxD3 (24). While the FoxD3 mutations in these cases were not the primary cause of the ASD phenotypes, it is believed that they increase the general risk of ASD phenotypes.

sox10, a member of the SOX family of transcription factors, is critical for the specification, and migration of NCCs throughout the body, laying the foundation for numerous tissues, organs, and cell types during the early stages of development (3, 10, 26, 27). sox10 is also a member of the expansive Gene Regulatory Network of NCCs and is essential for the formation of anything that is derived from the NCC including, but not limited to, blood vessels, enteric nerves, melanocytes, craniofacial cartilage and the inner ear (9, 10, 26). Cells expressing sox10 are also found in the AS (28). Initially, they are exclusive to the ventral regions of the eye, entering the AS via the choroid fissure as invading endothelial cells, and subsequently throughout the AS (1, 11, 13). Despite a critical role in NCC specification and migration, a functional role for sox10 in AS development has yet to be established.

Using fluorescent *in situ* hybridization as well as functional studies, we observed that the earliest ASM can be observed at 16hpf and exhibit tfap2a and foxc1a expression, but not that of foxd3 or sox10. Genetic knockout of either tfap2a or foxd3, but not sox10, results in the complete absence of ASM specification and contributes to malformation of the AS. We therefore conclude that the initial specification of ASM involves a tfap2a/foxc1a double positive subpopulation of cNCCs established soon after the optic cup if formed.

## RESULTS

### Timing of cranial neural crest specification into anterior segment mesenchyme

Since numerous studies in various vertebrate models point to the cranial neural crest (cNCC) as the origin of the anterior segment mesenchyme (ASM), we sought out to characterize this event in the zebrafish model. To do so, we first tracked the expression of canonical markers of the cNCC during early development (12hpf) up to the expected timing of initial ASM arrival to the eye (24hpf). The markers chosen were *sox10, foxD3* and *tfap2a*. Using two color fluorescent whole mount in situ hybridization (FWISH) we first compared expression of *foxd3* and *sox10* (Fig 1A-G). From as early as 12hpf *foxd3* and *sox10* show significant overlap in their signal confirming they are co-expressed by the migrating cNCC. The overlap is most apparent in the newly delaminating cNCCs found along the notochord and continues throughout the time course. By 20hpf we observe both *foxd3* and *sox10* expression surrounding the forming retina (periocular), but even at 24hp *foxd3* and *sox10* expression is absent from the anterior segment space and therefore ASM. When comparing expression of *tfap2a* and *sox10* we also observe near complete co-expression in the notochord and the periocular regions by 24hpf. However, in contrast to *sox10* or *foxd3*, at 16hpf we detect *tfap2a* positive cells surrounding developing retina as well as within the anterior segment space (Fig 1H-N). Interestingly, the tfap2a positive cells found surrounding the retina and within the AS space are negative for *sox10* or *foxd3* expression. In fact, a significant cranial ventral population of *tfap2a* positive but *sox10/foxD3* negative cells can be distinguished as early as 18hpf (Fig 1K). Tfap2a positive cells that first enter the anterior segment space, 16-18hpf, appear to do so as two streams, one dorsal one ventral (Fig 1L). Also, both streams appear to be restricted from the future lens area. By 24hpf, *tfap2a* positive cells are found through the AS space including the lens region. Having detected tfap2a positive cells in the AS space as early as 16hpf, much earlier than expected, we next examined if these cells were in fact early ASM. To do so we compared co-expression of *tfap2a* with well-established ASM markers *foxc1a, foxc1b* and *pitx2* (Fig 2). When comparing expression patterns of *tfap2a* and *foxc1a* we detected a high degree of co-expression in regions surrounding the retina as well as within the AS starting at 18hpf (Fig 2A-G). Interestingly, prior to 16hpf, *foxc1a* and *tfap2a* expression is spatially restricted with *foxc1a* concentrating in the developing forebrain and tfap2a concentrating in the notochord and tail regions. Co-expression of *foxc1a* and *tfap2a* in the AS was highly evident by 24hpf. Similar patterns were observed when comparing *tfap2a* and *foxc1b* expression, however the *foxc1b* signal is not as strong in the ASM as *foxc1a* at earlier timepoints (Fig 2H-N). When examining *pitx2* expression we noted a very distinct ventro-nasal expression at 12hpf, (Fig 2O-S) which persists and expands. The *pitx2* signal was detected in the AS by 24hpf (Fig 2U). *Pitx2* AS expression does coincide with *tfap2a* positive cells, however, the number of co-expressing cells is very low and concentrated to the dorsal rim of the eye. In fact, a majority of ASM *tfap2a* positive cells are *pitx2* negative at 24hpf. Lastly, we also compared *foxc1a* and *pitx2* expression to that of *sox10*. When comparing *foxc1a* with *sox10* we observed a high degree of co-expression along the notochord, starting as early as 14hpf and continuing up to 18hpf (Fig 3 A-G). We also noted a distinct population of *foxc1a* positive, *sox10* negative cells in the forebrain (Fig 3D). As expected, we did not detect any *sox10* and *foxc1a* double positive cells in the AS segment prior to 24hpf. All double positive cells were restricted to the dorsal rim of the retina. Similar outcomes were observed when comparing *pitx2* and *sox10* expression with only a few *pitx2/sox10* double positive cells observed in the very dorsal AS space. From our spatiotemporal analysis we concluded that the primary source of zebrafish ASM is derived from *tfap2a/foxc1a*, but not *sox10* or *foxd3* expressing cNCCs which arrive to at the AS space as early as 16hpf and represent the earliest ASM progenitors.

**Figure 1:**
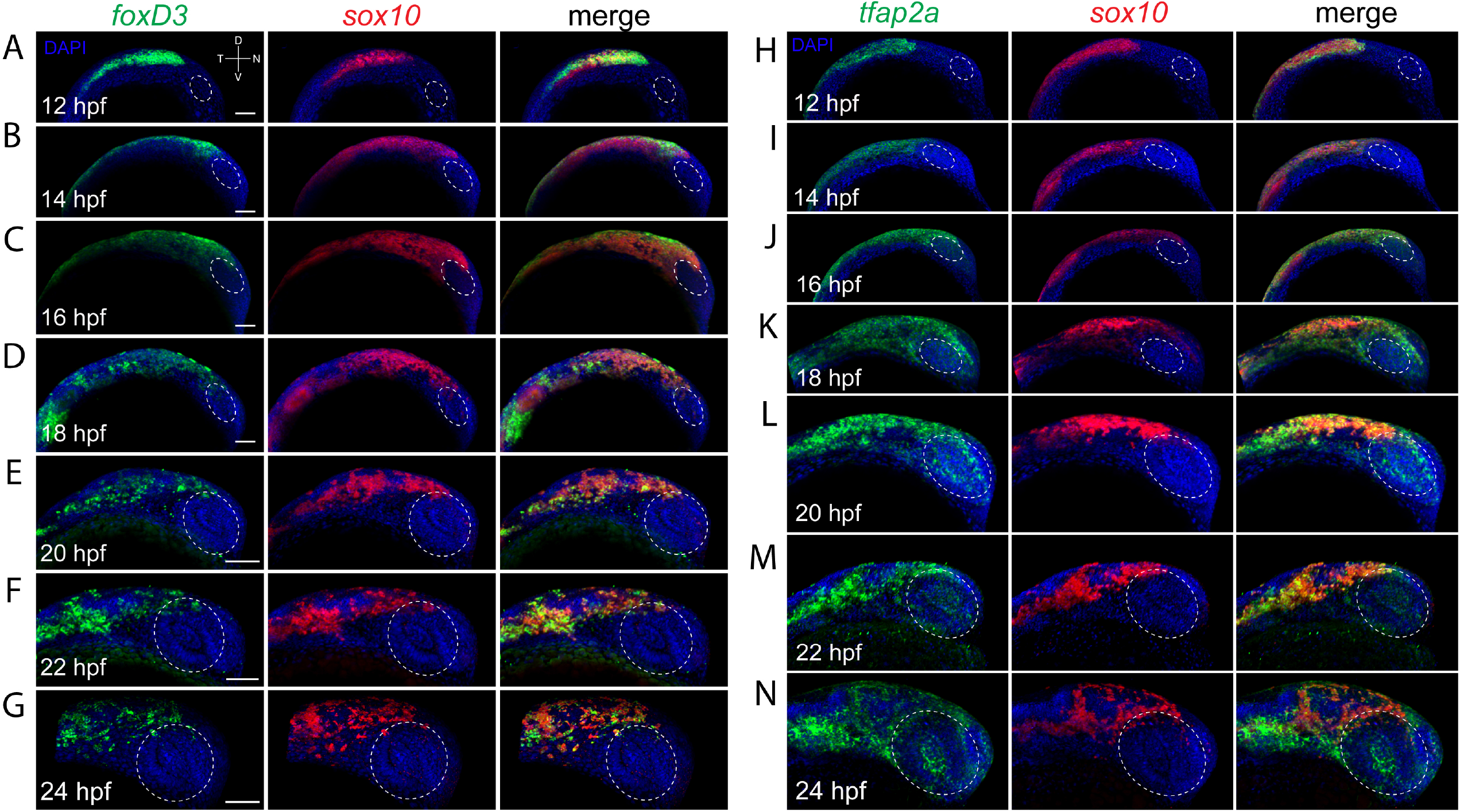
Co-expression analysis of cNCC markers during eye field and retinal morphogenesis. **A-G)** Two color FWISH analysis of *foxD3* (green) and *sox10* (red) at 12-24hpf with samples taken at 2h intervals. DNA was stained with DAPI (blue). Lateral images of volume projections of 3D confocal stacks from representative embryos are displayed. Strong co-expression of *foxD3* and *sox10* can be observed in the region of the notochord at early timepoints and throughout the neural crest stream starting at 16hpf. Periocular co-expression (white oval) is observed starting at 22hpf. **H-N)** Two color FWISH analysis of *tfap2a* (green) and *sox10* (red) at 12-24hpf with samples taken at 2h intervals. DNA was stained with DAPI (blue). Lateral images of volume projections of 3D confocal stacks from representative embryos are displayed. Strong co-expression of *tfap2a* and *sox10* can be observed in the region of the notochord as early as 12hpf and throughout the time course observed. Starting at 16hpf, *tfap2a* positive cells (yellow *) can be observed in the anterior segment (white oval). Scale bar = 100μm

**Figure 2:**
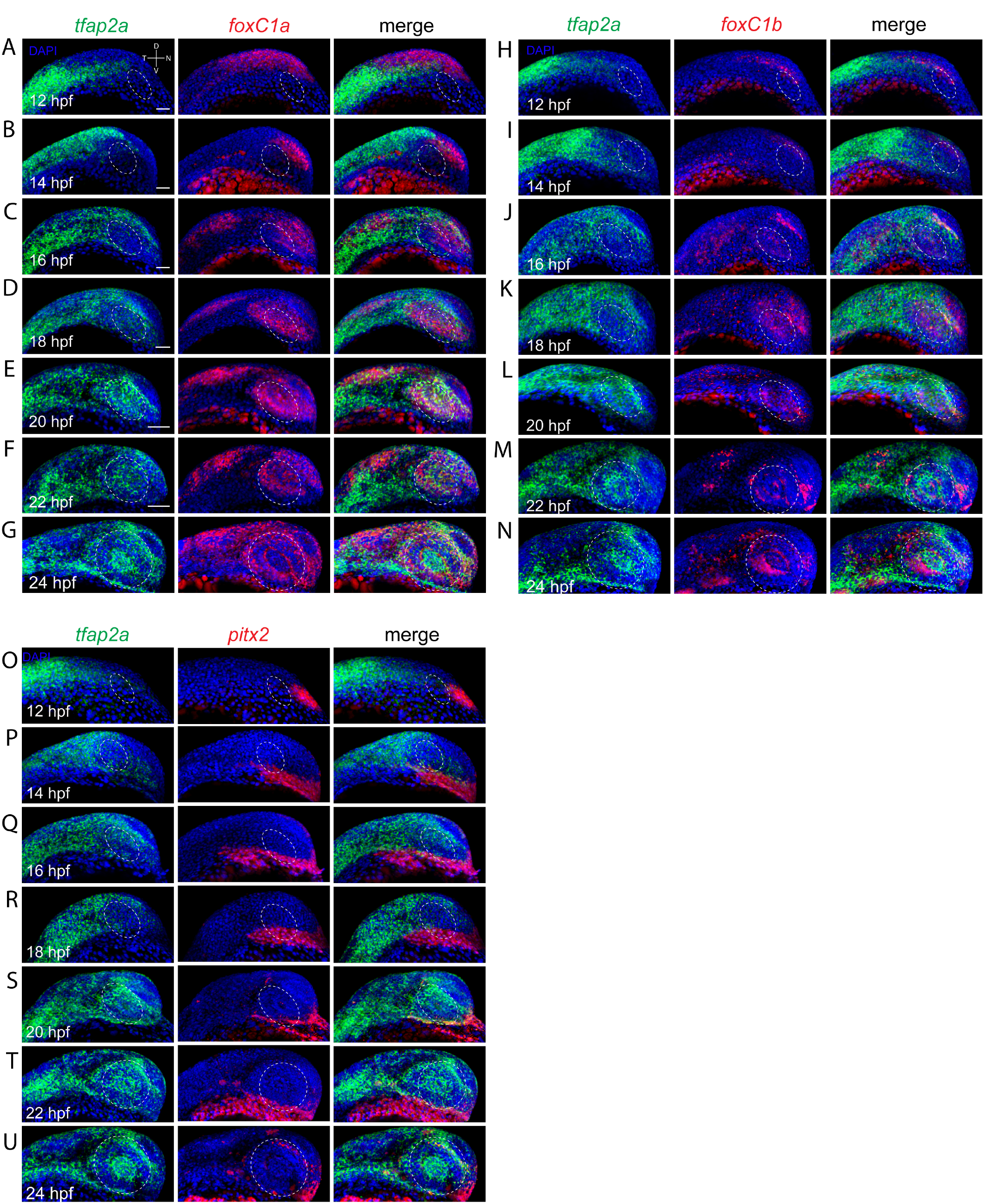
Co-expression analysis of *tfap2a* and anterior segment mesenchyme markers during eye field and retinal morphogenesis. **A-G)** Two color FWISH analysis of *tfap2a* (green) and *foxc1a* (red) at 12-24hpf with samples taken at 2h intervals. DNA was stained with DAPI (blue). Lateral images of volume projections of 3D confocal stacks from representative embryos are displayed. Co-expression of *tfap2a* and *foxc1a* first observed in the notochord and periocular regions at 16hpf. Starting at 18hpf, and extending to 24hpf, *tfap2a*/*foxc1a* double positive cells are detected in the anterior segment region (white oval). **H-N)** Two color FWISH analysis of *tfap2a* (green) and *foxc1b* (red) at 12-24hpf with samples taken at 2h intervals. DNA was stained with DAPI (blue). Lateral images of volume projections of 3D confocal stacks from representative embryos are displayed. Co-expression of *tfap2a* and *foxc1b* is first observed in the dorsal periocular regions (white oval) at 16hpf. Starting at 18hpf, and extending to 24hpf, *tfap2a/foxc1b* double positive cells are detected in the anterior segment region (white oval). **O-U)** Two color FWISH analysis of *tfap2a* (green) and *pitx2* (red) at 12-24hpf with samples taken at 2h intervals. DNA was stained with DAPI (blue). Lateral images of volume projections of 3D confocal stacks from representative embryos are displayed. Co-expression of tfap2a and *pitx2* is first observed in the ventral periocular regions at 20hpf. Co-expression of tfap2a and *pitx2* in the anterior segment (white oval) is observed starting at 24hpf in the very dorsal and ventral regions. Scale bar = 100μm

**Figure 3:**
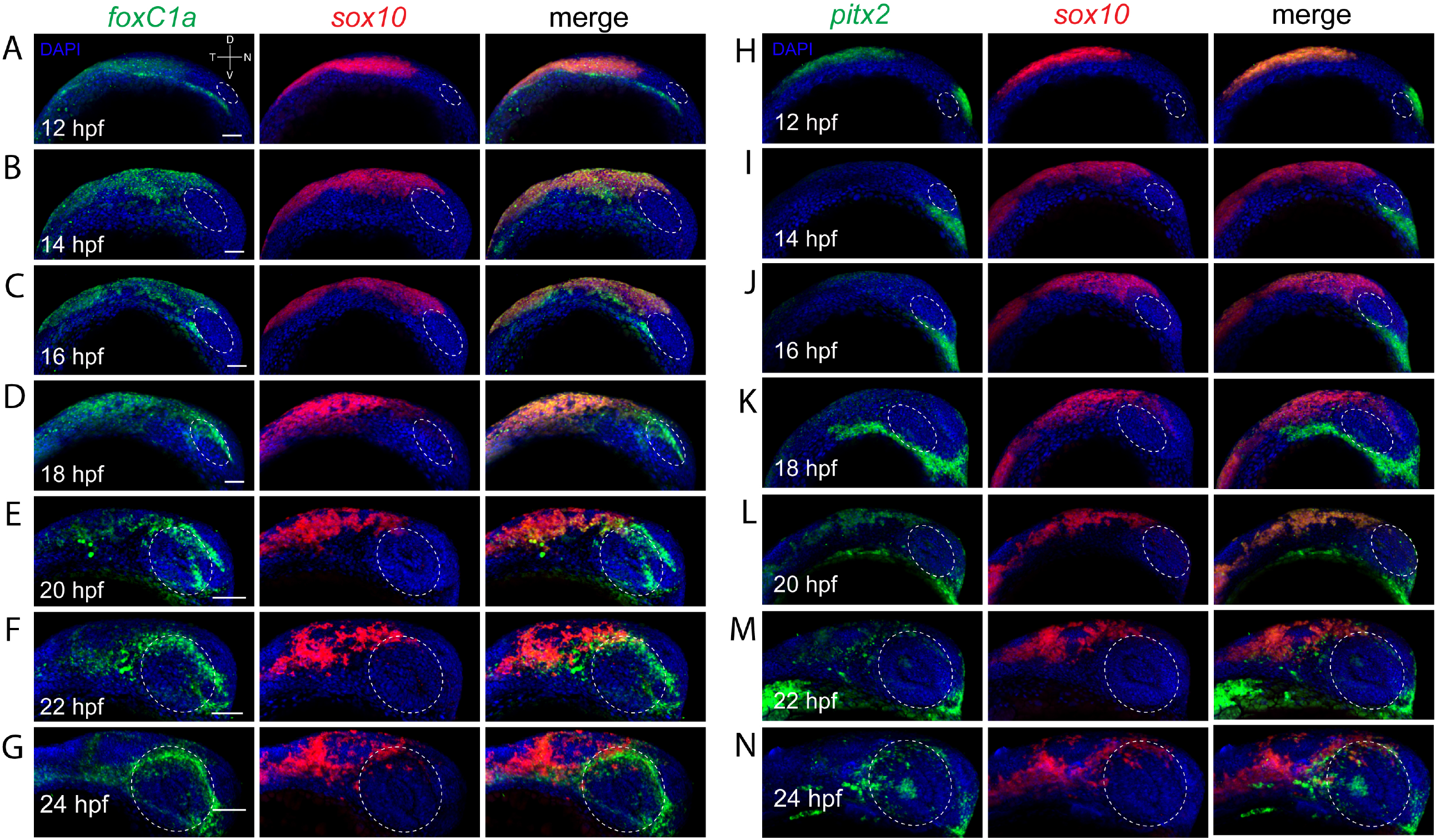
Co-expression analysis of *sox10* and anterior segment mesenchyme markers during eye field and retinal morphogenesis. **A-G)** Two color FWISH analysis of *foxc1a* (green) and *sox10* (red) at 12-24hpf with samples taken at 2h intervals. DNA was stained with DAPI (blue). Lateral images of volume projections of 3D confocal stacks from representative embryos are displayed. Strong co-expression between *sox10* and *foxc1a* is observed starting at 14hpf in the notochord. Anterior segment (white oval) expression for *foxc1a* is detected starting at 18hpf and continuing up to 24hpf while *sox10* expression is excluded from the anterior segment region in all timepoints. **H-N)** Two color FWISH analysis of *pitx2* (green) and *sox10* (red) at 12-24hpf with samples taken at 2h intervals. DNA was stained with DAPI (blue). Lateral images of volume projections of 3D confocal stacks from representative embryos are displayed. Co-expression between *sox10* and *pitx2* is first observed starting at 20hpf in the notochord but overall co-expression is minor. *Pitx2* expression is primarily constrained to regions ventral of the developing eye. By 22-24hpf *pitx2* expression is detected in the anterior segment space (white oval) and includes several cells exhibiting co-expression with sox10. Scale bar = 100μm

### Cranial neural crest specification into anterior segment mesenchyme requires function of tfap2a and foxd3

Having determined that tfap2a, but not sox10 or foxd3, is a marker the earliest zebrafish ASM progenitors, we next examined the consequences of the loss of tfap2a on ASM specification. To do so we employed the tfap2a^sa2445^ mutant line. This line harbors a single nucleotide substitution, A>T, resulting in a non-sense mutation in exon 4 at amino acid 236. Based on its location, the mutant allele results in a truncation of 201 amino acids and is therefore expected to generate a null phenotype. To examine ASM development in the absence of tfap2a function we collected clutches of embryos from tfap2a^+/sa2445^ in-crosses at 32hpf and assayed for the expression of ASM markers *foxc1a, lmx1b*.*1* and *eya2*. Based on previous results we expect robust labeling of ASM using these markers at 32hpf (1). Using FWISH, expected WT and tfap2a^+/sa2445^ embryos, approximately 75%, displayed very distinct AS expression patterns for *foxc1a, lmx1b*.*1* and *eya2*. All three exhibited expression along the periphery of the eye (periocular) as well as within the AS space, classical ASM expression patterns. Conversely, in approximately 25% of embryos, the expected ratio of tfap2a^sa2445/sa2445^, *foxC1a, lmx1b*.*1* and *eya2* expression was nearly absent (Fig 4A). In confirmed tfap2a^sa2445^ homozygous embryos we observe a total loss of both periocular and ASM expression for *foxc1a* and *lmx1b*.*1*, and significant reduction in expression of *eya2*, especially in the periocular regions. These results clearly point to the necessity of tfap2a function during specification of ASM.

**Figure 4:**
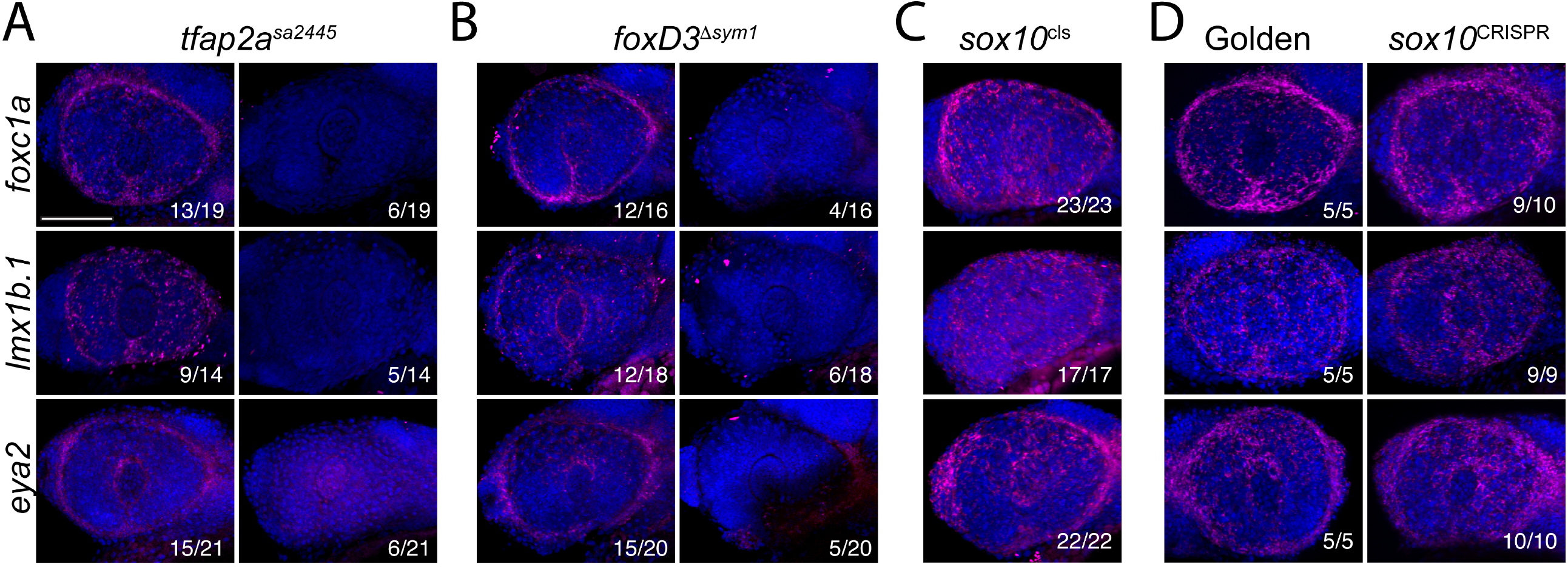
tfap2a and foxd3 are necessary for ASM specification. **A)** Fluorescent WISH analysis of *foxc1a, lmx1b*.*1* and *eya2* expression (magenta) in 32hpf embryos from tfap2a^sa2445/+^ in-crosses. DNA was stained with DAPI (blue). Lateral images of volume projections of 3D confocal stacks from representative embryos are displayed. Proportion of embryos with corresponding expression phenotypes are displayed. In approximately 25% of embryos *foxc1a, lmx1b*.*1* and *eya2* expression was severely reduced. **B)** Fluorescent WISH analysis of *foxc1a, lmx1b*.*1* and *eya2* expression (magenta) in 32hpf embryos from foxd3^sym1/+^ in-crosses. DNA was stained with DAPI (blue). Lateral images of volume projections of 3D confocal stacks from representative embryos are displayed. Proportion of embryos with corresponding expression phenotypes are displayed. In approximately 25% of embryos *foxc1a, lmx1b*.*1* and *eya2* expression was severely reduced. **C)** Fluorescent WISH analysis of *foxc1a, lmx1b*.*1* and *eya2* expression (magenta) in 32hpf embryos from sox10^cls/+^ in-crosses. DNA was stained with DAPI (blue). Lateral images of volume projections of 3D confocal stacks from representative embryos are displayed. Proportion of embryos with corresponding expression phenotypes are displayed. All embryos examined displayed expected, wildtype, expression phenotypes. **D)** Fluorescent WISH analysis of *foxc1a, lmx1b*.*1* and *eya2* expression (magenta) in 32hpf embryos injected with golden crRNA or sox10 crRNA (CRISPR). DNA was stained with DAPI (blue). Lateral images of volume projections of 3D confocal stacks from representative embryos are displayed. Proportion of embryos with corresponding expression phenotypes are displayed. In both, golden and sox10 crRNA injected embryos expression of *foxc1a, lmx1b*.*1* and *eya2* was not affected. Scale bar = 100μm.

Having confirmed the necessity of tfap2a function in ASM specification, we next sought to examine whether foxd3 or sox10 also play a role in ASM development. Tfap2a and foxd3 are both known regulators of cNCCs development and share regulatory duties. As such, despite the absence of foxd3 expression in early ASM, we felt warranted to also examine the consequences of foxd3 loss of function on ASM specification. To do so we used the foxd3^sym1^ line which harbors a single nucleotide deletion leading to a premature truncation within the winged helix domain and culminating in a null allele (23). Since we saw little co-expression of *foxd3* in the early ASM we anticipated little to no effect on ASM gene expression. However, when examining foxD3^sym1^ homozygous mutants we observed results mirroring what was observed for tfap2a^sa2445^ mutant embryos. Expression of both *foxc1a* and *lmx1b*.*1* was absent from the ASM of foxd3^sym1^ homozygous embryos while *eya2* expression was again severely downregulated (Fig 4B). This unexpected result likely stems from the fact that loss of foxd3 function has been shown to affect *tfap2a* expression at early developmental time points (23). As such, loss of foxd3 function likely leads to reduced expression of tfap2a and therefore leads to mis-regulation of ASM specification.

### sox10 function is dispensable for ASM specification and development

Having observed the effects on ASM specification in the absence of foxd3 function, despite *foxd3* expression being absent from early ASM, we also examined the effects on ASM development caused by the absence of sox10 function. To do so, we used the well-studied colorless line (sox10^cls^) which harbors a 1.4kb transposon insertion and results in a non-sense mutation with a significant truncation in the protein and therefore a likely null allele (29). The *cls* line has a distinct phenotype of having no pigment cells and therefore remaining transparent during development. When examining 32hpf embryos from clutches of sox10^cls/+^ heterozygous in-crosses, we observed wildtype like expression for all three of our ASM probes in all the embryos imaged. This result suggested that unlike tfap2a and foxd3, sox10 function is not necessary for ASM development and/or specification. To confirm the *cls* results, we also knocked out sox10 function using CRISPR-Cas9. Using the Alt-R-CRISPR platform from IDTDNA, we injected two adjoining crRNAs to target sox10 coding sequence and induce an approximate 2140bp deletion between exon 2 and exon 3. This large deletion results in a frameshift and subsequent nonsense mutation resulting in a null allele. Genomic DNA from sox10 double crRNA injected embryos were assayed for the presence of the deletion and over 90% of the injected embryos displayed the results of bi-allelic cutting. Furthermore, injected embryos displayed the colorless phenotype at 6dpf (absence of pigmentation), indicating that CRISPR injections were representative of sox10 loss of function. To examine ASM specification, sox10 double crRNA injected embryos were subjected to FWISH probing for *foxc1a, lmx1b*.*1* and *eya2* expression. As a control, we also examined embryos injected with the Golden control crRNA which targets a non-related intron sequence (30). All the injected embryos, including Golden control and double sox10 crRNA, displayed wildtype patterns of *foxc1a, lmx1b*.*1* and *eya2* expression. As such, the CRISPR results reaffirmed that sox10 function is not necessary for ASM specification.

Considering that expression of both sox10 and foxd3 is absent from the earliest ASM, yet loss of foxd3 function has a consequence on ASM specification, we also wanted to eliminate the possibility of developmental delay in the foxD3^sym1^ line. As such, we examined eya2 and pitx2 expression in at 48hpf in foxd3^sym1^, sox10^cls^ and as an additional control the pitx2^sny15^ line. The sny15 line harbors a 10bp insertion resulting in a frame shift and premature stop codon in the second helix of the pitx2 homeodomain, therefore generating a null allele of pitx2. Loss of pitx2 function is known to result in a lack of ASM specification/development and therefore maldevelopment of the AS. As expected, in both pitx2^sny15^ and foxD3^sym1^ homozygous embryos *eya2* and *pitx2* expression was virtually lost, while all embryos from the sox10^cls^ line displayed wildtype expression. As such, we conclude that foxD3 but not sox10 is required for ASM specification despite not being expressed in the early tfap2a positive ASM.

Lastly, to determine whether absence of tfap2a or foxD3 function, and therefore specification of ASM, results in AS maldevelopment we examined AS structure in homozygous mutant larva at 6dpf. We identified 6dpf larva homozygous for tfap2a^sa2445^, foxD3^sym1^, sox10^cls^ as well as sox10 double crRNA injected larva and examined the AS in whole mounts as well as retinal cryosections stained with toluene blue (Fig 5C). Whole mount images of 6dpf tfap2a^sa2445/sa2445^ and foxD3^sym1/sym1^ both display abnormal AS morphology along with malformed and mispositioned eyes when compared to WT (Fig 5C). Tfap2a^sa2445/sa2445^ larva also display pigmentation defects. When comparing AS morphology in cryosections both tfap2a^sa2445/sa2445^ and foxD3^sym1/sym1^ larva have much smaller ASs, including thinning corneas and malformed ICAs. In contrast, sox10^cls/cls^ and CRISPR injected larva display a wild type like AS morphology in whole mounts as well as cryosections. As noted previously sox10^cls/cls^ and CRISPR injected larva display a complete lack of pigment in the head region, confirming sox10 loss of function. Corneas and ICAs in these larvae look highly like wild type controls. Taken together, we confirm that upon disruption of ASM specification via functional inactivation of tfap2a or foxd3 but not sox10, the AS does not develop properly due to improper specification of the ASM.

**Figure 5:**
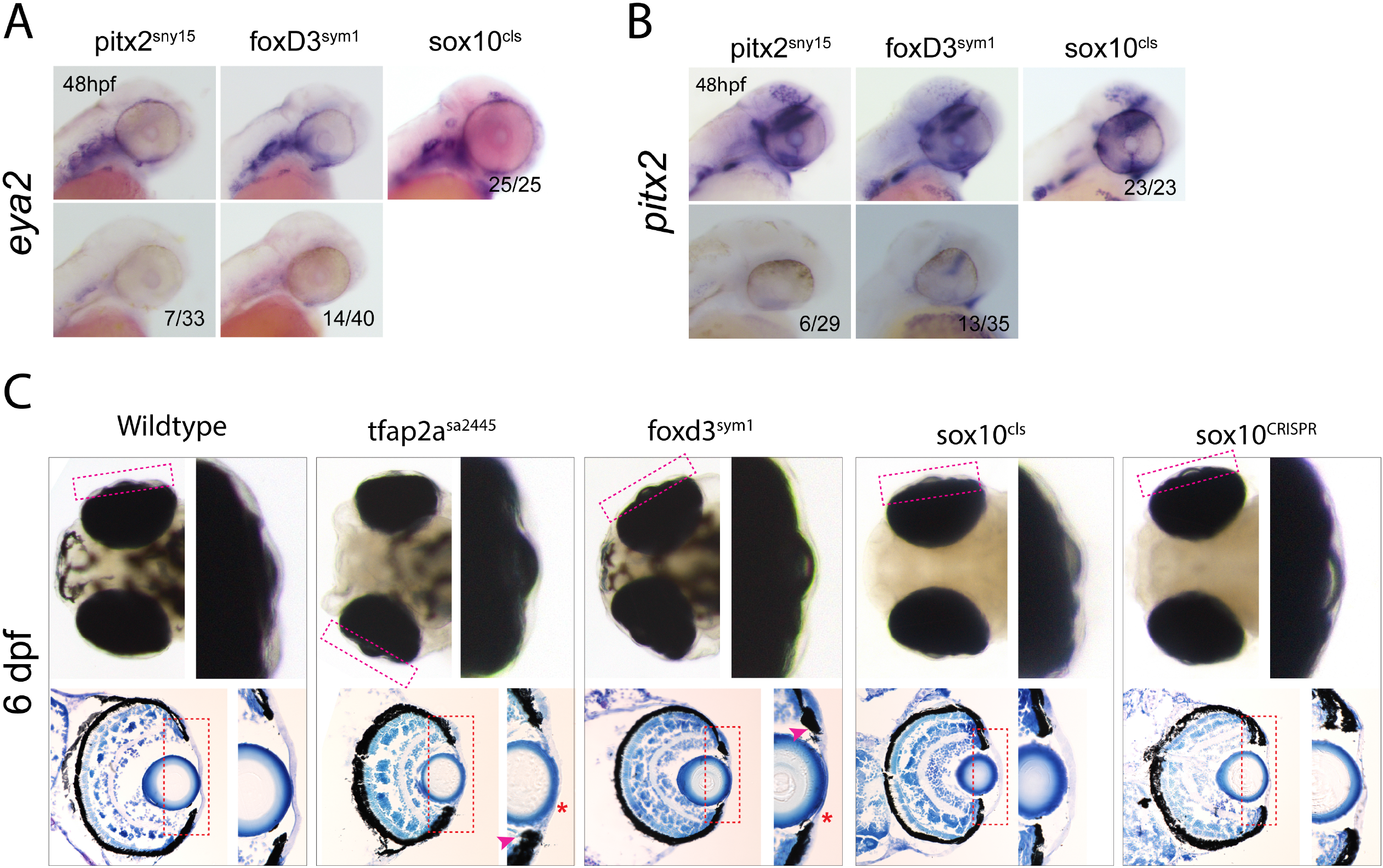
Anterior segment development does not require sox10 function. **A)**WISH detecting expression of *eya2* in 48hpf embryos from in-crosses of pitx2^sny15/+^, foxd3^sym1/+^ or sox10^cls/+^. Proportion of embryos with corresponding expression phenotypes are displayed. Approximately 25% of embryos from pitx2^sny15/+^ or foxd3^sym1/+^ in-crosses display the absence of eya2 expression, while 100% of embryos from sox10^cls/+^ crosses show wildtype levels of expression at 48hpf. **B)** WISH detecting expression of *pitx2* in 48hpf embryos from in-crosses of pitx2^sny15/+^, foxd3^sym1/+^ or sox10^cls/+^. Proportion of embryos with corresponding expression phenotypes are displayed. Approximately 25% of embryos from pitx2^sny15/+^ or foxd3^sym1/+^ in-crosses display the absence of *pitx2* expression, while 100% of embryos from sox10^cls/+^ crosses show wildtype levels of expression at 48hpf. **C)** Whole mount dorsal images and cryosections of wildtype, tfap2a^sa2445/sa2445^, foxd3^sym1/sym1^, sox10^cls/cls^ or sox10^CRISPR^. Cranial structures, improperly set eyes, as well as the anterior segment (Magenta rectangle representing the enlargement) thickness and organization are malformed in tfap2a^sa2445/sa2445^, foxd3^sym1/sym1^ larva but not in sox10^cls/cls^ or sox10^CRISPR^. Toluidine blue staining of 6dpf old eyes reveals a thinned cornea (red *) and irregular shape of the ICA (magenta arrow) in tfap2a^sa2445/sa2445^ and foxd3^sym1/sym1^ larvae, but not in sox10^cls/cls^ or sox10_CRISPR_.

### Transcriptomic analysis confirms divergent relationship between sox10 derived cNCC and POM

Having observed significant differences in spatiotemporal expression patterns and functional contribution of sox10 during ASM specification we lastly sought to compare transcriptomic profiles of foxd3, foxc1b and sox10 derived cNCC and POM. We hypothesized that foxd3 lineage derived cNCC/POM should contain POM and ASM transcriptomic profiles similar to that of foxc1b derived cells while sox10 lineages should significantly differ since sox10 has no functional consequence on POM/ASM specification. To test our hypothesis, we used transgenic reporter lines Tg[*sox10*:GFP], Tg[*foxc1b*:GFP] and Tg[*foxd3*:GFP] with the goal of isolating cNCC/POM/ASM populations and subsequently performing total RNA sequencing. We isolated heads of 48hpf Tg[*sox10*:GFP], Tg[*foxc1b*:GFP] or Tg[*foxd3*:GFP] embryos, therefore isolating cNCCs, POM and ASM cells, dissociated the tissue to make a single cell suspension and FACS sorted for GFP+ cells. At 48hpf ASM are firmly specified and have populated the AS space making this time points appropriate to determine if sox10 derived cells contribute significantly to the ASM cell fate. Approximately 500,000+ cells from each reporter line were collected in three individual replicates and subsequently sequenced the isolated total RNA. On average 40+ million reads were generated from each of the samples. Sequencing files processed to enable analysis and comparisons of expression patterns between the data sets. First, when using principal component analysis (PCA) to compare the three data sets Tg[*foxc1b*:GFP] and Tg[*foxd3*:GFP] samples displayed much closer association to each other, and therefore relatedness, than either of them to that of Tg[*sox10*:GFP] (Fig 6A). This supported our findings that foxd3 but not sox10 function was critical for ASM specification and therefore AS development. Second, when analyzing individual gene expression between our three samples we again noted clear clustering between the Tg[*foxc1b*:GFP] and Tg[*foxd3*:GFP] samples but not Tg[*sox10*:GFP] (Fig 6B). Lastly, gene ontology enrichment comparisons indicated that the sox10 isolated cells exhibited profiles of pigment, vasculature and muscle function while having an absence of keratinization and other AS related expression patterns found in the Tg[*foxd3*:GFP] and Tg[*foxc1b*:GFP] cells (Fig 6C). Taken together, transcriptomic analysis supports the hypothesis that despite sox10 being a key NCC regulator, sox10 lineage derived cNCC have a minor contribution to ASM and AS development.

**Figure 6:**
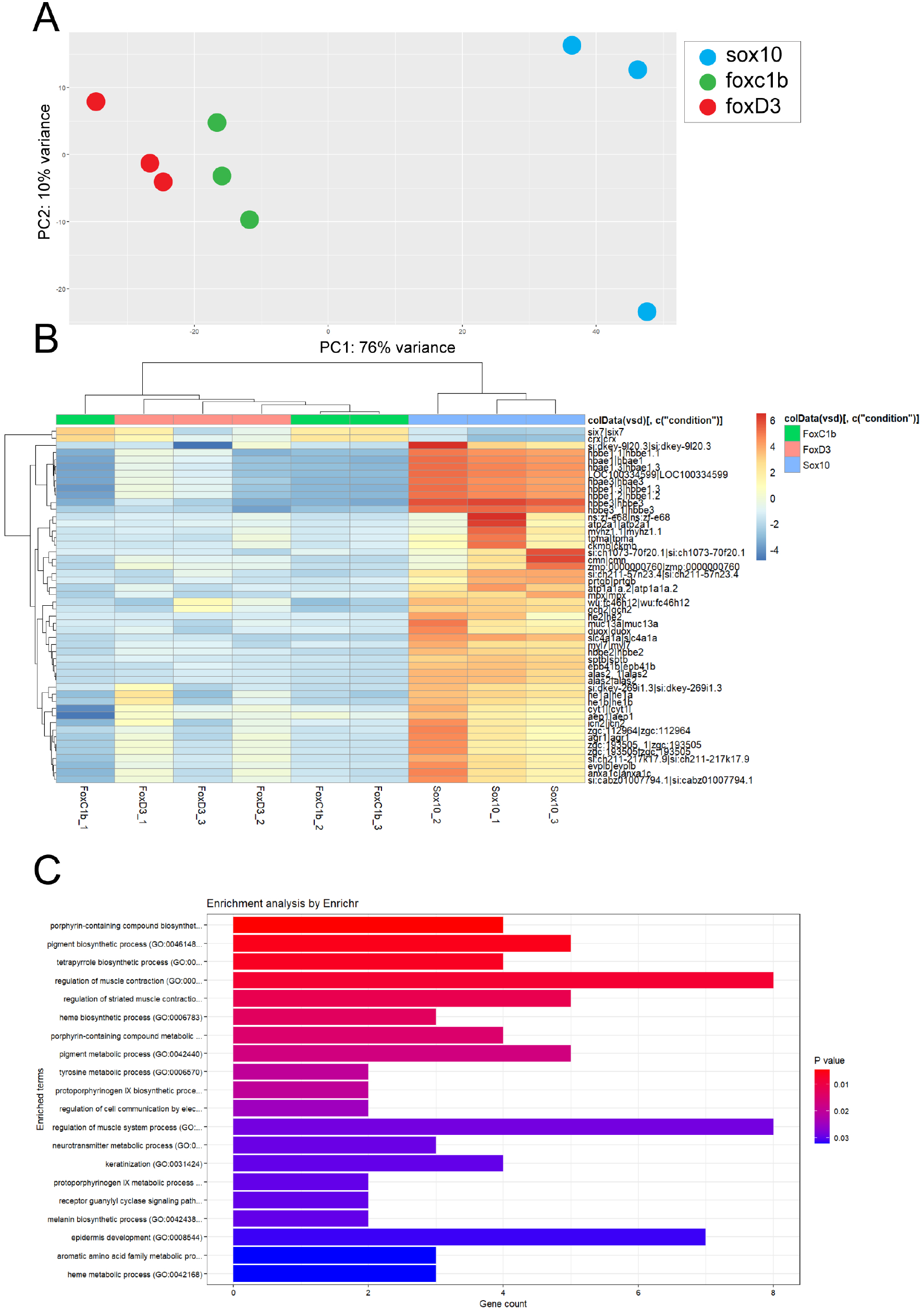
Transcriptomic comparison between sox10, foxd3 and foxc1b derived cNCCs. **A)** Principal component analysis comparing 48hpf Tg[*sox10*:GFP] (blue), Tg[*foxd3*:GFP] (red) and Tg[*sox10*:GFP] (green) isolated cNCC/POM transcriptomes. Each dot represents one biological replicate. Variance between the samples is least between foxd3 and foxc1b data sets which cluster together and far apart from sox10 samples. **B)** Cluster analysis of the biological replicates indicates a close relationship between foxd3 and foxc1b derived samples and a significant divergence from sox10 derived samples. Numerous genes are found to be differentially expressed between the foxd3/foxc1b and sox10 samples. **C)** Ontology enrichment analysis identifies sox10 datasets to exhibit an upregulation in pigmentation, muscle and hematopoietic biological pathways compared to ASM related processes such as keratinization.

## DISCUSSION

The NCC has been a topic of scientific inquiry for decades. This uniquely vertebrate population of cells has been associated with numerous functions and developmental consequences, including the contribution of cNCC to the formation of the anterior segment. However, while considerable effort has been made to identify and characterize the timing and regulatory mechanisms for specification of NCC into fully differentiated tissues, we still largely lack a detailed understanding of how cNCC are ultimately specified into ASM. In our current study we aimed to begin answering that question by tracking the expression patterns and functional contribution of critical cNCC regulators; tfap2a, foxd3 and sox10. Using fluorescent in situ hybridization, we observed that in zebrafish cNCC begin to populate AS as early as 16-18hpf. We made this conclusion based upon detecting co-expression of both ASM and cNCC associated genes *foxc1a* and *tfap2a* respectively in the anterior segment space as early as 16hpf. We clearly show that the earliest ASM found in the AS space are *foxc1a* and *tfap2a* double positive and arrive there between 16-18hpf. Surprisingly, we did not detect the expression of either *foxd3* or *sox10* in these early ASM. In fact, neither *sox10* nor *foxd3* expression is detected in ASM until about 24hpf, and even then, it is present in only very few cells. This suggests that a subpopulation of cNCCs which become tfap2a/foxc1a positive, no longer rely on foxd3 or sox10 function. We previously found by examining transgenic lines labeling various populations of ASM, including foxd3 and sox10 lineage derived cells, that the colonization of the ASM begins at 24-26hpf and proceeds in a dorsal to ventral pattern (1). Our current study suggests that those later arriving ASM cells may comprise a second wave of already specified cNCCs, while the initial wave includes the *tfap2a/foxc1a* positive cNCCs arriving as early as 16hpf. A colonization in multiple waves of the AS has been documented in mammalian species and our data now suggest it may also be conserved in the teleost linage (31). Our study also uncovers another interesting and possibly overlooked aspect of ASM specification. The expression of *foxc1a* in the forebrain region of the early embryo (14-18hpf) does not coincide with *tfap2a* expression. This finding suggests that a population of foxc1a positive cells exists independent of NCCs but may contribute to the earliest ASM. Future investigation into this cell population warrants effort as it may reveal a novel source of ASM progenitors.

In addition to observing the highly dynamic expression patterns associated with tfap2a, foxd3 and sox10 we also concluded that tfap2a and foxd3, but not sox10 function is critical for ASM specification and ultimately AS development. Genetic knockout of tfap2a or foxd3 both resulted in an almost complete absence of ASM marker gene expression at 32hpf (Fig 4) as well as corresponding defects in the AS at 6dpf (Fig 5). Thinning corneas and disorganized ICAs were observed at 6dpf in tfap2a^sa2445^ and foxd3^sym1^ mutants. These results were in stark contrast to both the sox10^cls^ mutant or the sox10^CRISPR^ induced loss of function where larvae from both displayed little to no effect on ASM specification or AS formation (Fig 5). Ultimately, these results confirmed our hypothesis where tfap2a regulates ASM specification, but at the same time offered surprising new insights. First, due to its lack of expression in early ASM, foxd3 was expected to have little to no effect on ASM or AS development. However, foxd3^sym1^ mutants clearly show a drastic reduction in ASM marker gene expression and AS deficiencies at 6dpf (Figs 4-5). We attribute this outcome to the fact that foxd3 is a key regulator of not only cNCCs but NCCs in general. As such, its loss likely leads to a general reduction or misspecification of cNCC, therefore affecting those early ASM. In support of this hypothesis, it’s been shown that foxd3 mutations can result in misexpression of tfap2a (23). Second, sox10 is also a key regulator of cNCCs and NCCs, and yet its absence had no effect on ASM or the development of the AS. Both the sox10^cls^ mutant line and our CRISPR induced loss of function approach had little to no effects on ASM marker gene expression and AS formation (Figs 4-5). In both cases we did observe a complete lack of pigmentation, confirming the loss of sox10 function. Hence, while foxd3 indirectly affected ASM and the AS, sox10 did not, despite being considered a critical regulator of the cNCC. To further examine this finding, we also analyzed expression levels in periocular mesenchyme cells, which also include ASM cells. These cells are derived from transgenic lines representing the sox10, foxd3 and foxc1b lineages. The goal was to determine whether lineages expected to contribute to the ASM, such as foxc1b, were similar or dissimilar to those from cNCC populations, foxd3 or sox10. RNA sequencing analysis revealed a close association between foxc1b and foxd3 cells, supporting the findings of our functional data. Though, sox10 cells did not share expression patterns associated with either foxc1b or foxd3. Ontology enrichment analysis revealed that both foxc1b and foxd3 lineages express marker genes associated with AS formation, such as migratory processes and keratogenesis. On the other hand, sox10 cells displayed enrichment in processes such as pigmentation, muscle formation and vasculogenesis. Based on the transcriptomic analysis, we can conclude that the functional differences we observed between tfap2a/foxd3 and sox10 can be explained by the differential nature of sox10 lineages of NCC when compared to foxc1b or foxd3. The sox10 associated or derived NCCs are not significant contributors to ASM or AS formation. In previous studies, sox10 was shown to be expressed in the ASM between 24-48hpf (1, 32). As such, sox10 expression in ASM may be a remnant of the general specification of NCC and in zebrafish but appears to have no significant functional role in subsequent ASM mediated AS development.

In conclusion, we have shown that ASM begin to assemble the anterior segment as early as 16hpf in zebrafish and that the major regulators of this process are tfap2a and foxd3. Future studies will focus on the identity and molecular signatures of these early ASM, as well as tracking their migratory patterns and behaviors.

## MATERIALS AND METHODS

### Zebrafish maintenance

Zebrafish lines were bred and maintained in accordance with IACUC regulations at the University of Kentucky (protocol # 2015-1370). AB strain was used as wildtype. sox10^cls^ was obtained from ZIRC, foxd3^sym1^ and tfap2a^sa2445^ lines were a gift from Dr. Lister. Transgenic lines used, Tg[sox10:GFP, foxc1b:GFP and foxd3:GFP] were described previously (1). All embryos were raised for the first 22 hours post fertilization in embryo media (E3) at 28°C. Unless indicated, after 22 hours, E3 media was replaced with embryo media containing 1-phenyl 2-thiourea (PTU) every 24 hours to maintain embryo transparency.

### *In situ* hybridization

Whole mount in situ hybridization (WISH) and Fluorescent WISH (FWISH) were performed as previously described (1). WISH and FWISH probes were generated using primers with T7 promoter sequences as previously described (1). List of primers used is found in table 1. WISH embryos were imaged using a Nikon Di2 5MP camera mounted on the Nikon SMZ800 stereo scope using Nikon Elements software and processed for figure presentation using Adobe photoshop and Illustrator. FWISH stained embryos were mounted in low melting point agarose using glass bottom dishes (Fluoro dish) and imaged using the C2+ Nikon confocal microscope using a 20X NA 0.95 objective. 3D images were collected at 2.5um intervals. Images were processed using Adobe photoshop and illustrator software.

### Alt-R-CRISPR injections

crRNA, trcRNA and Cas9 was purchased from IDTDNA. Duplexing of crRNA with trcRNA was performed as previously described (33). Two duplexed crRNAs were co-injected with Cas9 to generate an approximately 2140bp deletion between exon 2 and exon 3 of sox10. crRNA targeting sequences were: CAGCCACGTCGACGCAAGAA and CACCCCCAAGACGGAACTGC. Injection parameters used were as described previously (33). Confirmation of Cas9 cutting was done using PCR with primers flaking the cut sites by 150bp up and down stream (Table 1). gDNA isolated from embryos was used to assess efficiency of cutting.

### Cryosections

6dpf embryos, grown without PTU, were euthanized using MS222 and subsequently fixed in 4% paraformaldehyde for 24 hours. Post fixation embryos were treated in 10% and subsequently 30% sucrose for 24hours each. Post sucrose washes the embryos were mounted in OCT (optimal cutting temperature) embedding medium and frozen in cryosection molds. Cryosections were collected at 10μm thickness using a Leica cryotome. Sections were subsequently air dried for 24h and stained with 1% toluidine blue for 2min at room temperature and washed with ddH20 and two 95% and 100% EtOH washes followed by three xylene washes. Slides were then mounted using a xylene based mounting medium (cytoseal). Images were collected using a Nikon Ti2 compound microscope with a 20X NA 0.95 objective. Subsequent processing was performed using Adobe photoshop and illustrator.

### RNA sequencing

Collected Tg[*foxd3*:GFP], Tg[*foxc1b*:GFP] or Tg[*sox10*:GFP] embryos were incubated in E3 media at 28°C up to 48hpf. At this time, embryos were dechorionated, anesthetized using 3-amino benzoic acidethylester (Tricaine) and decapitated just posterior of the eyes. Heads were collected and incubated for 2 minutes in 0.25% Trypsin + EDTA at 37°C. After incubation, a 20G needle and syringe were used to dissociate the tissue before the tube was placed back at 37°C for 2min. This process was repeated 4 times. After incubation, the dissociated cells were strained using a 40μm filter (VWR) and spun down for 10mins at 3500rpm at 4°C. The supernatant was removed, and the pellet resuspended in 1x PBS + 2 mM EDTA. Cells were sorted for GFP+ cells at the University of Kentucky Flow Cytometry and Immune Monitoring Core at the Markey Cancer Center. ∼250,000-500,000 cells were collected in three distinct trials. Following cell sorting, RNA was extracted using Trizol according to the manufacturers protocol. Concentration and purity of extracted RNA was performed using the Bioanalyzer at the University of Kentucky HealthCare Genomics Core at the UK Albert B. Chandler Hospital. RNA was then DNAse treated (TURBO DNAfree Kit, Thermo Fisher) and sent to Applied Biological Materials (ABM) for Illumina sequencing. 20-40 million 150bp paired-end reads were generated from each of the 3 samples/transgenic line examined. Fastq files for each sample were annotated using the NCBI reference genome using hisat2, then converted into bam files using stringtie. Gtf files were then used to make a gene count matrix which were analyzed by R. DEseq2 was used to calculate principal component analysis and generate gene expression heat maps. Gene ontology enrichment was completed using Enrichr.

## ACKNOWLEDGEMENTS

We thank Dr. James Lister for his generosity with transgenic and mutant zebrafish lines. We also thank Dr. Jeramiah Smith for his assistance with bioinformatics analysis of RNAseq data and Dr. Ann Morris for use of the cryo-microtome. Lastly, we acknowledge the excellent zebrafish husbandry work of Brandi Bolton. This work was supported by the NIH-NEI grant R01EY027805-01A1.

---

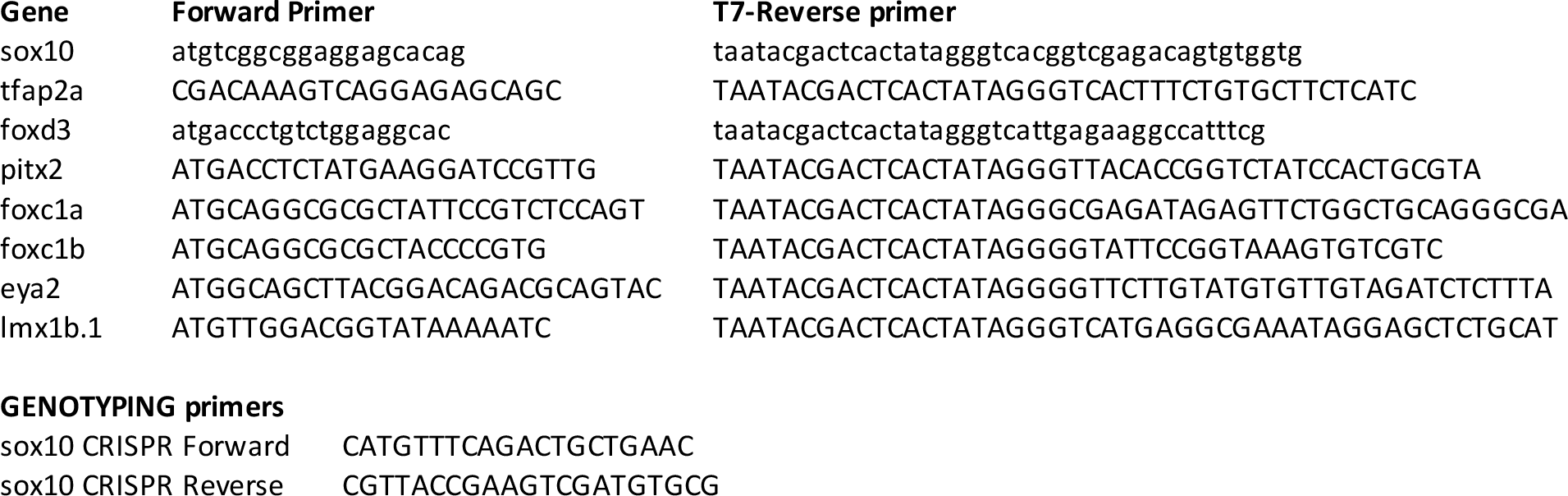
IN SITU HYBRIDIZATION probe PRIMERS.

